# A Two-Stage ESM-Based Machine Learning Pipeline for Robust Hierarchical Enzyme Function Prediction

**DOI:** 10.64898/2026.08.14.744831

**Authors:** Xiao Hua, Ghjuvan M. Grimaud

## Abstract

Accurate enzyme annotation remains a major bottleneck in translating rapidly growing protein sequence data into biological knowledge. Enzyme Commission (EC) prediction is particularly challenging because enzyme functions are organized hierarchically, annotations are often imbalanced across classes, and sequence similarity alone may be insufficient to resolve functional differences. To address these challenges, we developed ESM-ECForest, a two-stage framework that combines protein embeddings generated by the pretrained language model ESM-2 (Evolutionary Scale Modeling 2) with Random Forest classifiers. The first stage distinguishes enzymes from non-enzymes, whereas the second assigns one or more EC numbers to proteins predicted to be enzymatic. On an external benchmark comprising 25,778 protein sequences, ESM-ECForest achieved the highest weighted F1 score among the evaluated methods at all four EC levels, decreasing from 0.94 at Level 1 to 0.90 at Level 4. The largest relative improvements were observed for lyases (EC 4), ligases (EC 6), and translocases (EC 7), although EC 6 and EC 7 remained the most difficult classes internally. Visualization of the ESM-2 embedding space using Uniform Manifold Approximation and Projection (UMAP) revealed clustering patterns consistent with enzyme functional relationships, indicating that biologically relevant information is retained in the pretrained representations prior to supervised classification. These results support the use of pretrained protein language model embeddings as an effective foundation for enzyme annotation. By combining large-scale sequence representations with a lightweight supervised classifier, ESM-ECForest provides a scalable approach for EC prediction and may facilitate functional annotation of protein sequences derived from large genomic and metagenomic datasets.

## 1 Introduction

The rapid growth of genome and metagenome sequencing has led to an unprecedented accumulation of protein sequence data. In contrast, experimentally supported functional annotations remain available for only a small fraction of known proteins. This imbalance is particularly evident in microbial datasets, where many predicted proteins show limited similarity to characterized enzymes and therefore remain difficult to annotate reliably. Because enzymes connect gene products to specific biochemical reactions, accurate enzyme annotation plays a central role in interpreting cellular functions and reconstructing metabolic processes [1,2].

The Enzyme Commission (EC) system provides a standardized framework for describing enzyme function through a hierarchical classification scheme that reflects the biochemical reactions catalyzed by enzymes [3–5]. Beyond protein function annotation, EC numbers are widely used in pathway reconstruction, metabolic network analysis, and genome-scale metabolic modelling, where they serve as an important link between protein sequences and biochemical reactions, including their equations and stoichiometry.

Experimental characterization remains the most reliable approach for enzyme annotation [4]. However, functional validation is often constrained by the availability of suitable substrates, cofactors, and assay conditions, making large-scale characterization impractical [2]. As a result, computational approaches have become indispensable for narrowing the gap between sequence availability and functional knowledge. A wide range of computational strategies has been developed for enzyme annotation [6–9], including sequence similarity-based methods [10,11], homology-driven approaches [12,13], structure-informed prediction [14,15] and machine learning (ML) frameworks [16–25]. Sequence alignment tools such as BLAST continue to be widely used because of their simplicity and interpretability. Their effectiveness, however, decreases as sequence similarity declines, particularly when distant homologues possess distinct biochemical functions [26–28]. This challenge has motivated increasing interest in machine-learning approaches that learn sequence–function relationships directly from annotated datasets.

Recent models such as DeepEC [20], ProteInfer [24] and DeepECTransformer [21] have demonstrated that enzyme function can be inferred from amino acid sequences using supervised learning strategies. Nevertheless, the performance of these approaches remains influenced by the characteristics of the available training data. Enzyme annotations are unevenly distributed across EC classes, with well-studied enzyme families represented by substantially more examples than rare or poorly characterized activities [9,29]. Such imbalances complicate model training and can limit predictive performance for underrepresented enzyme classes and newly sequenced proteins [18,20].

Protein language models (PLMs) provide an alternative approach to protein sequence representation. Models such as ESM-2 are pre-trained on large collections of protein sequences using self-supervised objectives, enabling them to learn contextual representations without relying on functional annotations [29–31]. Previous studies have shown that these representations capture information related to protein structure, evolutionary relationships, and biological function, supporting a variety of downstream applications including structure prediction and functional inference. Unlike conventional supervised approaches, PLMs can exploit information contained in large unlabeled sequence collections, potentially reducing dependence on annotated training datasets [27,32,33]. Despite these advances, the extent to which pretrained protein representations improve enzyme annotation across the EC hierarchy remains unclear [34]. In particular, classes with limited annotation coverage, substantial functional diversity, or low sequence conservation continue to pose challenges for existing prediction methods [16,29,35–38]. Understanding whether information encoded by protein language models can improve robustness in these settings is therefore of considerable practical interest.

In the present study, we developed ESM-ECForest, a two-stage framework that combines ESM-2-derived protein embeddings with Random Forest classifiers for enzyme identification and EC prediction. The first stage distinguishes enzymes from non-enzymes, whereas the second performs hierarchical EC assignment for predicted enzymes. By separating sequence representation learning from downstream classification, the framework allows information learned from large-scale protein sequence corpora to be incorporated into enzyme annotation. We evaluated ESM-ECForest using homology-aware data partitions and independent benchmark datasets and compared its performance with established enzyme annotation tools, including DeepEC, ProteInfer and DeepECTransformer. The results provide an opportunity to assess the contribution of pretrained protein representations to hierarchical enzyme annotation and to examine their utility in large-scale functional analysis.

## 2 Methods

### 2.1 ESM-ECForest: two-stage identify-then-classify workflow

The ESM-ECForest workflow is summarized in Fig. 1. Protein sequences were first converted into fixed-length numerical representations using ESM-2. The resulting embeddings served as input features for all downstream classifiers. In stage A, a binary Random Forest classifier was trained to distinguish enzymes from non-enzymes. Sequences predicted as enzymatic were subsequently passed to stage B. In stage B, enzyme sequences are assigned one or more EC numbers through a multi-label classification framework. This two-stage design separates enzyme detection from functional classification, allowing EC prediction to be restricted to proteins predicted to possess catalytic activity.

**Fig. 1.**
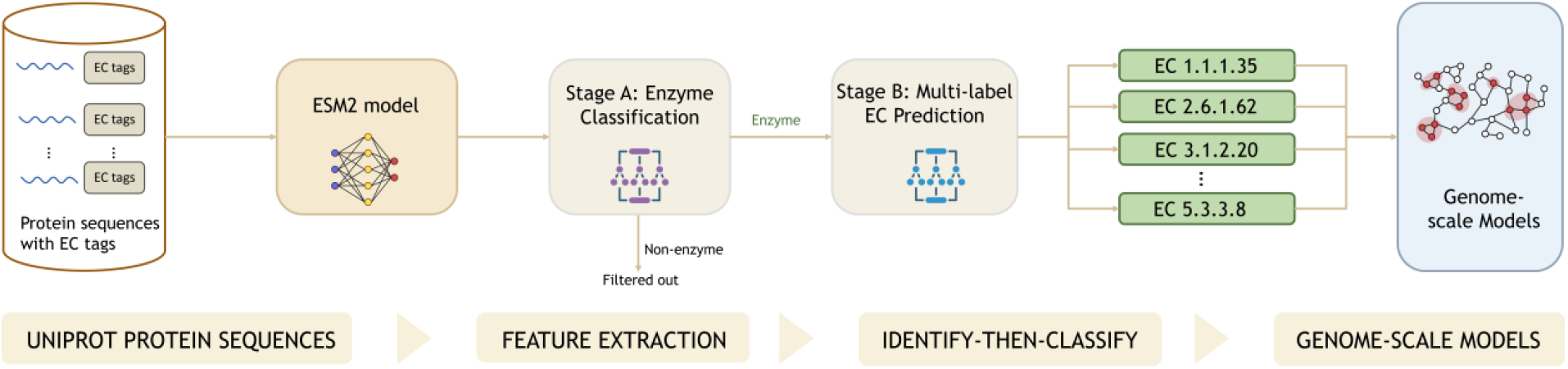
Overview of the ESM-ECForest workflow. Protein sequences obtained from UniProt were transformed into fixed-length embeddings using ESM-2. Stage A performed enzyme/non-enzyme classification, and proteins predicted as enzymes were subsequently evaluated by the stage B classifier for hierarchical EC prediction. The resulting EC annotations can be linked to biochemical reactions and used in downstream functional analyses.

Model performance was assessed both across the complete EC hierarchy and within individual top-level EC classes. To facilitate comparison with existing approaches, ESM-ECForest was benchmarked against DeepEC, ProteInfer, and DeepECTransformer using an independent evaluation dataset. In addition, Uniform Manifold Approximation and Projection (UMAP) was used to visualize the ESM-2 embedding space and to examine the distribution of enzyme and non-enzyme and EC-class representations.

### 2.2 Dataset construction for ESM-ECForest training and validation

Protein sequences and functional annotations were obtained from UniProtKB/Swiss-Prot (release 2026_01) and used to construct datasets for enzyme identification and EC number prediction. The dataset was assembled from representative prokaryotic and unicellular eukaryotic taxa selected to maximize phylogenetic diversity while limiting overrepresentation of a small number of model organisms. Inclusion of phylogenetically distant groups was intended to improve taxonomic coverage and support the evaluation of enzyme prediction across diverse regions of protein sequence space. The complete list of taxa included in the study is provided in Supplementary Table S1.

Data curation followed procedures similar to those adopted in previous enzyme annotation studies [18,22]. Only entries with UniProt annotation scores of 4 or 5 were retained. The annotation score reflects the completeness of sequence annotation and ranges from 1 to 5, with higher values indicating more extensively curated entries [39]. Sequences shorter than 50 amino acids or longer than 5,000 amino acids were excluded. In the collected datasets, sequences were classified as non-enzyme if the reported catalytic activity and EC annotations were absent, in line with [22]. For multifunctional enzymes, all available EC numbers were retained and assigned to the corresponding protein sequence, resulting in a multi-label annotation scheme. Among the 104,648 enzyme sequences in the curated dataset, 21,103 (20.2%) were annotated with multiple EC numbers and were classified as multifunctional enzymes. The final dataset consisted of 104,648 enzymes and 80,366 non-enzymes, corresponding to a total of 185,014 protein sequences.

To reduce sequence redundancy, all sequences were clustered using MMseqs2 (v17-b804f) at 40% sequence identity and 80% coverage. Dataset partitioning was performed at the cluster level with a target train/validation/test ratio of 8:1:1, ensuring that no cluster was shared between subsets. Owing to variation in cluster size, the resulting sequence counts deviated from the target ratio. Stratified sampling was applied whenever possible to preserve class distributions. For enzyme identification, stratification was based on enzyme versus non-enzyme labels, whereas EC classification used the primary EC annotation. Classes represented by fewer than two clusters were assigned randomly. Dataset statistics are provided in Supplementary Table S2.

### 2.3 Training and validation of ESM-ECForest

Training and model selection were performed using the cluster-based dataset partitions described above. The training set was used for model fitting, the validation set for hyperparameter optimization and threshold selection, and the test set for internal model evaluation during model development.

For stage A, the binary classifier was trained using enzyme and non-enzyme sequences from the training set. For stage B, only enzyme sequences were used, with EC annotations encoded as multi-label targets. Multifunctional enzymes contributed to all associated EC labels during training.

Hyperparameters were optimized on the validation set. The evaluated parameters included the number of trees, maximum tree depth, minimum samples required for node splitting, feature-sampling strategy, class-weight configuration, and prediction threshold. The final parameter set was selected according to validation-set performance measured by weighted F1 score.

Class imbalance in the binary enzyme identification task was addressed using balanced class weights. For EC prediction, label imbalance was accounted for during evaluation by reporting macro-, micro-, and support-weighted metrics. After hyperparameter optimization, the final model was fixed and subsequently evaluated using the independent benchmarking dataset described below (section 2.8). The benchmarking dataset was not used for model fitting, parameter tuning, threshold optimization, or model selection. All analyses were performed using a fixed random seed of 42 to ensure reproducibility.

### 2. Protein representation using ESM-2

Protein sequences were encoded using ESM-2, a pretrained protein language model trained on large-scale protein sequence data through a masked-token prediction objective [32]. The model generates context-dependent sequence representations from primary amino acid sequences.

In this study, sequence embeddings were generated using the esm2_t6_8M_UR50D model implemented in the FAIR ESM Python package (fair-esm v2.0.0). This model contains six transformer layers and approximately 8 million parameters.

For each sequence, the beginning-of-sequence (BOS) token representation from the final transformer layer was extracted as a fixed-length feature vector of 320 dimensions. For sequences longer than 1000 amino acids, non-overlapping sequence chunks were encoded independently and the resulting embeddings were averaged to obtain a single protein-level representation. These embeddings served as input features for all downstream classification tasks, including enzyme identification and EC prediction [32,38].

### 2.5 Random Forest classifiers

Random Forest was used as the downstream classifier for mapping ESM-2 embeddings to enzyme-related labels [40]. All classifiers were implemented in Python using *scikit-learn* (v1.6.1) and trained on the 320-dimensional ESM-2 embeddings described above.

In stage A, enzyme identification was formulated as a binary classification problem. A Random Forest classifier consisting of 400 trees (maximum depth = 40) was trained to distinguish enzymes from non-enzymes. Class imbalance was addressed using balanced class weights. Additional parameter settings are provided in Supplementary Table S3.

In stage B, EC annotation was formulated as a multi-label classification task and was performed only for proteins predicted as enzymes. EC annotations were encoded using MultiLabelBinarizer, and a One-vs-Rest strategy was adopted in which a separate binary Random Forest classifier was trained for each EC label. The training dataset contained 3,021 distinct EC labels, resulting in 3,021 individual classifiers. Multiple EC numbers could therefore be assigned to a single protein by predicting more than one positive label. During inference, each stage B classifier generated a probability score for its corresponding EC label. EC labels with predicted probabilities ≥ 0.7 were retained as final annotations. The threshold was selected based on validation-set performance and fixed before evaluation on the independent benchmark dataset. Random Forest was selected because it can accommodate high-dimensional feature representations without requiring additional feature engineering and has been widely applied to biological classification tasks [41,42]. Its ensemble-based structure also provides a practical approach for handling heterogeneous datasets and nonlinear relationships between protein representations and functional labels.

### 2.6 Evaluation metrics

Model performance was assessed using precision, recall, and F1 score. True positives (TP), false positives (FP), and false negatives (FN) were determined by comparing predicted labels with reference annotations [9,43].

For a given class:

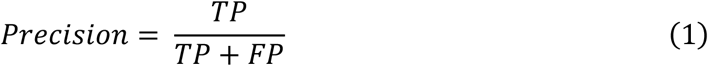

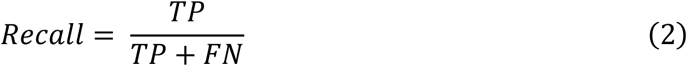

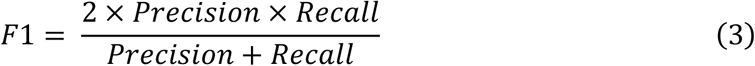

For the binary enzyme identification task, precision, recall, and F1 score were computed for enzyme versus non-enzyme classification. For EC number prediction, performance was evaluated in a hierarchical multi-label framework. Each EC annotation associated with a protein was treated as an independent positive label [44]. Missing annotated EC numbers were counted as false negatives, whereas unsupported additional EC predictions were counted as false positives. Predicted and reference EC numbers were compared at EC Levels 1-4 after truncation to the corresponding hierarchical level. For example, EC 1.2.3.4 contributed to the labels 1, 1.2, 1.2.3, and 1.2.3.4 at Levels 1-4 respectively. Duplicate labels generated during truncation were removed prior to metric calculation. Incomplete EC annotations (e.g., 1.2.-.-) were evaluated only at supported hierarchical levels and were excluded from deeper-level comparisons.

Hierarchical performance was evaluated independently at each EC level using precision, recall, and F1 score. Multi-label EC prediction performance was summarized using macro-, micro-, and support-weighted averages. At each EC level, label-specific precision (*P*_*i*_), recall (*R*_*i*_) and F1 score (*F*1_*i*_) were calculated according to Eqs. (1)–(3).

For *K* EC labels, macro-averaged metrics were calculated as:

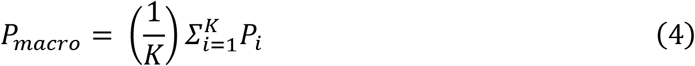

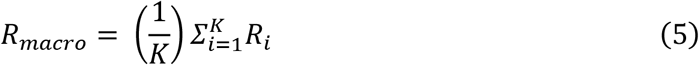

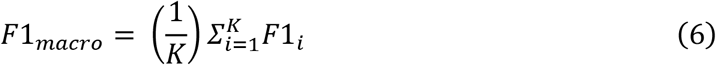

Micro-averaged metrics were calculated after pooling true positives, false positives, and false negatives across all labels:

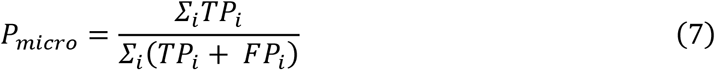

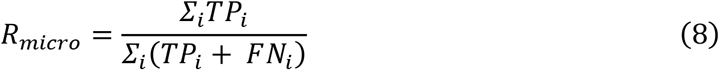

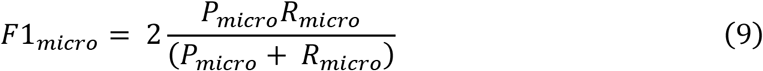

Support-weighted metrics were calculated as:

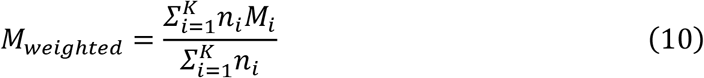

where *M*_*i*_ denotes the precision, recall, or F1 score for label *i*, and *n*_*i*_ represents the number of reference instances associated with that label.

Confidence intervals were estimated by bootstrap resampling of test-set proteins (1,000 replicates). Precision, recall, and F1 score were recalculated for each replicate, and 95% confidence intervals were defined by the 2.5th and 97.5th percentiles of the resulting distributions.

Performance was additionally evaluated for each first-level EC class, and class support was reported alongside the corresponding metrics. Proteins without predicted EC annotations were counted as false negatives for the corresponding reference labels. EC classes unsupported by a given method were retained in the evaluation and treated as missing predictions. Undefined precision or recall values were assigned a value of zero. The same evaluation procedure was applied to ESM-ECForest, DeepEC, ProteInfer, and DeepECTransformer.

### 2.7 UMAP visualization of embedding space

The 320-dimensional ESM-2 embeddings were visualized using Uniform Manifold Approximation and Projection (UMAP) [45]. Two-dimensional projections were generated for (i) the complete enzyme/non-enzyme dataset and (ii) enzyme sequences coloured by first-level EC class. Because the full dataset exceeded 180,000 sequences, a stratified subset of 30,000 proteins was used for visualization. Stratification preserved both enzyme/non-enzyme proportions and first-level EC class distributions.

### 2.8 Dataset used for benchmarking

An independent benchmarking dataset comprising 25,778 protein sequences from 15 prokaryotic and unicellular eukaryotic species was assembled to compare ESM-ECForest with existing enzyme annotation tools. The selected species represent diverse phylogenetic groups and include organisms commonly used in enzyme annotation and metabolic reconstruction studies. The complete species list is provided in Supplementary Table S4. Among the 14,847 proteins with EC annotations in the benchmarking dataset, 7,528 (50.7%) carried multiple EC numbers and were therefore considered multifunctional enzymes.

Benchmark proteins were not included in the training, validation, or internal test sets used for the development of ESM-ECForest. Sequence similarity between benchmark and training proteins was assessed using MMseqs2. The highest observed sequence identity was 38.4%, which remained below the 40% sequence identity threshold used for MMseqs2 clustering during dataset construction, indicating minimal sequence overlap between the benchmarking and training datasets.

### 2.9 Benchmarking against existing tools

ESM-ECForest was benchmarked against three sequence-based enzyme annotation tools: DeepEC, ProteInfer, and DeepECTransformer [20,21,24]. These methods were selected because they represent different modelling strategies for protein function prediction, including convolutional neural networks, dilated convolutional architectures, and transformer-based approaches.

DeepEC performs EC number prediction using convolutional neural networks supplemented by homology-based annotation for sequences that cannot be confidently classified by the neural network component[20]. ProteInfer predicts enzyme functions directly from unaligned amino acid sequences using a deep dilated convolutional architecture [24]. DeepECTransformer combines transformer-based sequence modelling with homology search and was developed for large-scale enzyme annotation, particularly in microbial genomes [21].

Benchmarking was performed using an external dataset described in section 2.8. Official implementations provided by the respective authors were used for all methods. Unless otherwise stated, analyses were performed using the recommended default settings. Repository commit hashes are reported in Supplementary Table S5 where formal software releases were unavailable. All predictions were mapped to a common EC-number representation before evaluation.

Because the evaluated methods differ in EC coverage and output format, all predictions were converted to a common EC representation before evaluation. Precision, recall, weighted F1 score, and per-class F1 score were calculated as described in Section 2.6. Incomplete EC annotations were evaluated only at the highest supported EC level. Unsupported classes were retained in the overall hierarchical evaluation as missing predictions, whereas unsupported class-specific values were displayed as N/A in per-class plots.

Runtime and peak memory usage were evaluated using the benchmark dataset described above. Wall-clock time and peak memory consumption were measured during prediction only, from execution of the prediction command to completion. Preprocessing steps unrelated to prediction, including software installation, database and model downloads, and output standardization, were excluded. Hardware and software specifications of the benchmarking environment are summarized in Supplementary Table S6.

## 3 Results

### 3.1 Performances of ESM-ECForest

#### 3.1.1 Enzyme identification performance of stage A

Stage A of ESM-ECForest was designed to distinguish enzymes from non-enzymes prior to EC annotation. On the independent benchmarking dataset (n = 25,778 proteins), the model achieved a precision of 0.975 (95% CI, 0.972-0.977), a recall of 0.977 (95% CI, 0.975-0.980), and an F1 score of 0.976 (95% CI, 0.974-0.978) (Fig. 2A). The complete performance results are provided in Supplementary Table S7.

**Fig. 2.**
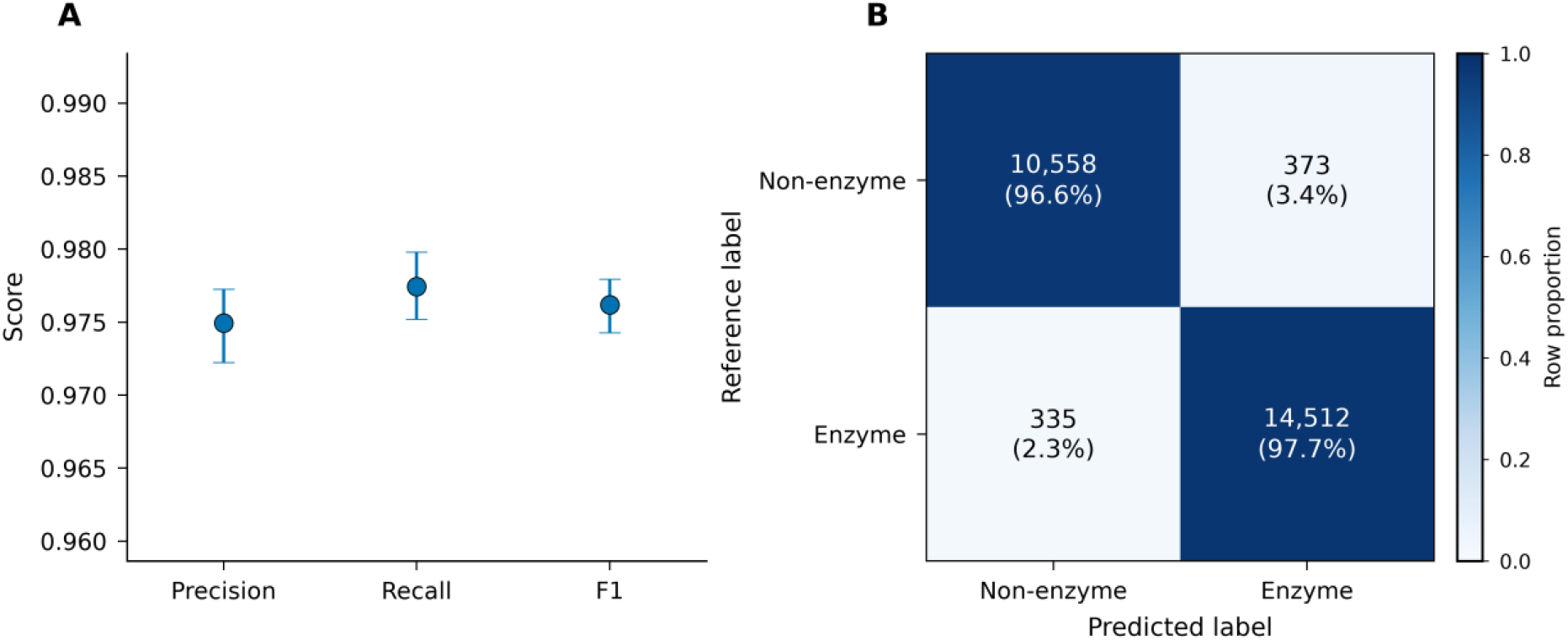
Binary enzyme identification performance of ESM-ECForest. (A) Precision, recall, and F1 score evaluated on the independent benchmarking dataset. Error bars denote bootstrap 95% confidence intervals. (B) Confusion matrix for enzyme and non-enzyme classification. Values indicate the number of proteins, with row-wise percentages shown in parentheses.

The corresponding confusion matrix revealed low error rates for both classes (Fig. 2B), with 373 non-enzymes incorrectly classified as enzymes and 335 enzymes classified as non-enzymes. As a result, 97.7% of enzymes and 96.6% of non-enzymes were correctly identified. Together, these findings indicate that Stage A achieves robust and well-balanced binary classification performance, providing a reliable foundation for subsequent EC number prediction.

#### 3.1.2 EC prediction performance of the complete ESM-ECForest framework

Following the evaluation of binary enzyme identification, we assessed the end-to-end performance of ESM-ECForest for hierarchical EC prediction across Levels 1-4 of the EC classification system (Fig. 3). As expected, predictive performance gradually decreased with increasing annotation depth under all averaging schemes (Fig. 3A-C). The decline was most pronounced for macro-averaged metrics, whereas micro- and weighted-average metrics remained comparatively stable across hierarchy levels.

**Fig. 3.**
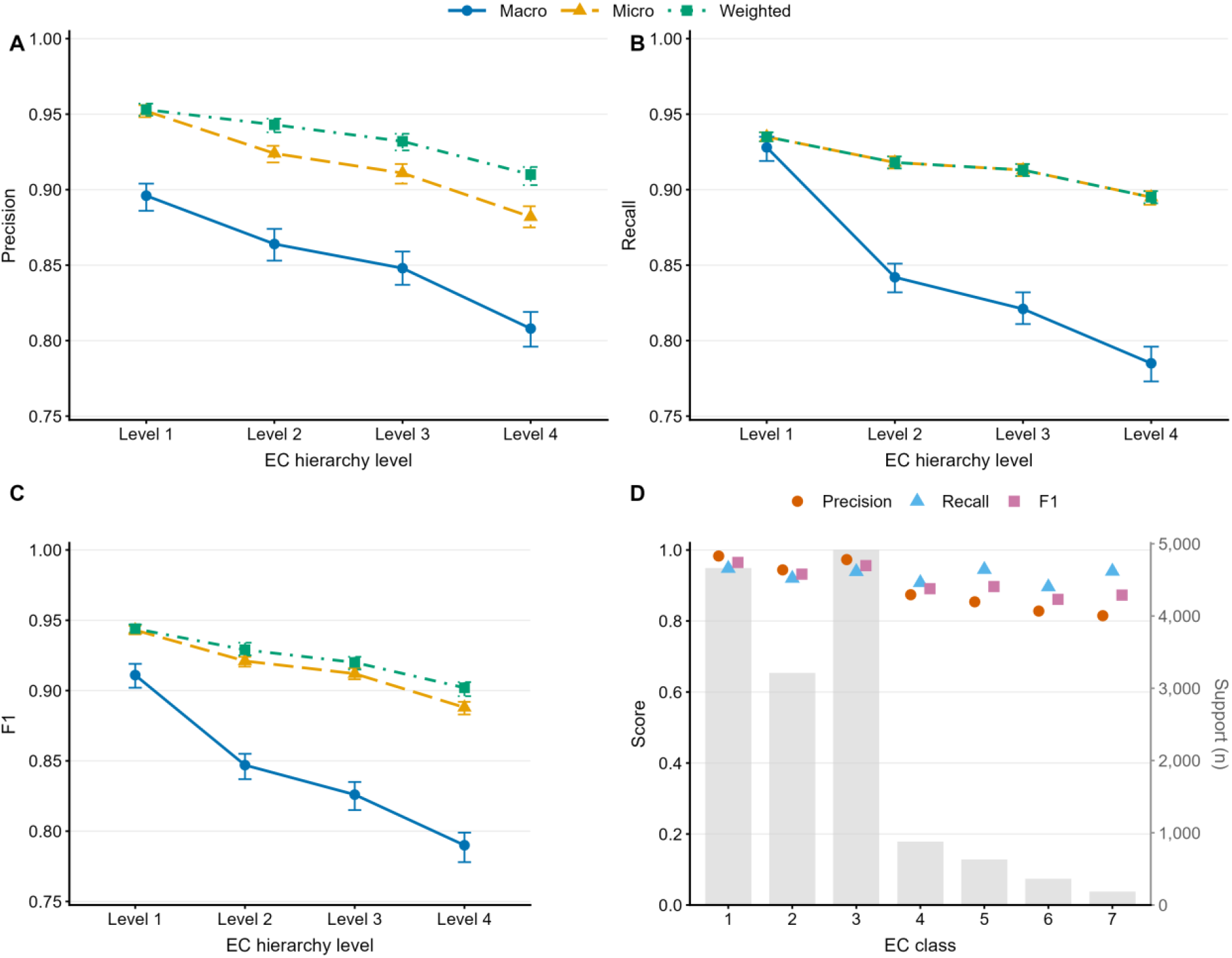
Hierarchical and class-specific EC prediction performance of ESM-ECForest. (A–C) Precision, recall, and F1 score across EC hierarchy Levels 1–4 using macro-, micro-, and support-weighted averaging. Points indicate performance estimates on the independent benchmarking dataset, and error bars denote 95% confidence intervals obtained from 1,000 bootstrap resamples. (D) Precision, recall, and F1 score for individual first-digit EC classes at Level 1. Grey bars indicate the number of reference annotations for each class (support; right y-axis).

At EC Level 1, macro-, micro-, and weighted-average F1 scores were 0.911 (95% CI, 0.902-0.919), 0.943 (95% CI, 0.940-0.946), and 0.944 (95% CI, 0.941-0.947), respectively. At Level 4, these values decreased to 0.790 (95% CI, 0.778-0.799), 0.888 (95% CI, 0.883-0.892), and 0.902 (95% CI, 0.896-0.906). Similar trends were observed for precision and recall (Fig. 3A,B). Macro precision decreased from 0.896 to 0.808, while macro recall decreased from 0.928 to 0.785 between Levels 1 and 4. In contrast, weighted-average metrics showed only modest declines across hierarchy levels and remained substantially higher than the corresponding macro-averaged metrics. Detailed performance results are provided in Supplementary Table S8A.

To further examine class-specific behaviour, performance was evaluated separately for each first-level EC category (Fig. 3D). EC 1-3 achieved F1 scores of 0.965, 0.932, and 0.956, respectively, whereas EC 4-7 showed lower performance, with F1 scores ranging from 0.861 to 0.897. Precision and recall also varied across EC classes, indicating differences in classification difficulty among enzyme categories. Detailed class-specific results are provided in Supplementary Table S8B. Despite these differences, all first-level EC classes achieved F1 scores above 0.86, demonstrating robust prediction performance across the major enzyme categories. Per-EC F1 scores were compared with the log-transformed number of training proteins associated with each EC label (Supplementary Fig. S1). Performance varied widely across EC labels, particularly among low-frequency labels, and no clear relationship between training-set frequency and per-EC performance was observed. Proteins were also grouped according to their maximum sequence identity to the training dataset. At EC Level 4, both macro- and weighted-average F1 scores increased with sequence identity (Supplementary Fig. S2). Non-zero performance was observed across all sequence-identity ranges, including proteins without detectable matches in the training dataset.

#### 3.1.3 UMAP visualization of ESM-2 embedding space

To examine the organization of the pretrained protein representations, the 320-dimensional ESM-2 embeddings were projected into two dimensions using UMAP (Fig. 4). The projection containing both enzyme and non-enzyme sequences revealed partial separation between the two groups (Fig. 4A). Several regions were enriched for either enzymes or non-enzymes, although overlap between the two populations was also observed.

**Fig. 4.**
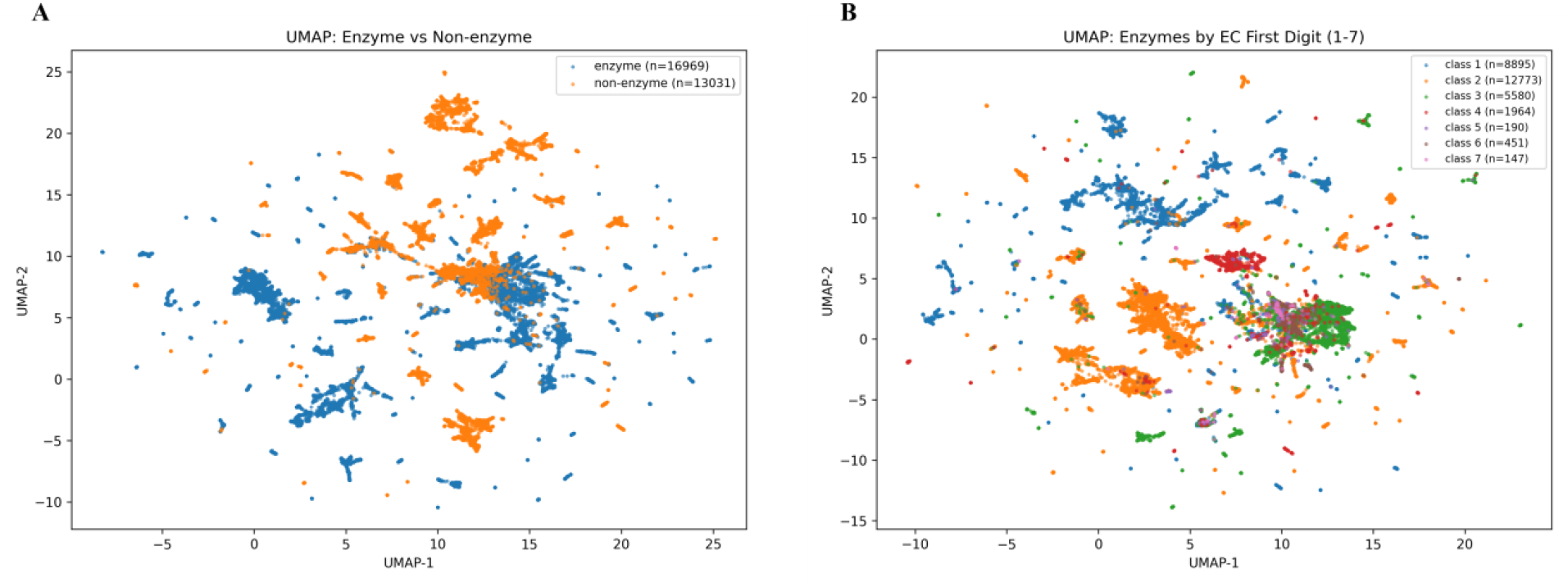
UMAP visualization of ESM-2 protein embeddings. (A) Projection of enzyme and non-enzyme sequences. (B) Projection of enzyme sequences colored according to first-level EC class.

A second UMAP projection was generated using enzyme sequences only and coloured according to first-level EC class (Fig. 4B). Distinct enrichment patterns were observed for several EC categories. The largest classes, including EC 1, EC 2, and EC 3, occupied broader regions of the embedding space, whereas EC 5, EC 6, and EC 7 were represented by smaller and more dispersed clusters. Considerable overlap remained among EC classes, particularly in regions containing multiple functional groups.

The UMAP projections indicate that proteins with different annotations are not uniformly distributed within the ESM-2 embedding space. Similarities in the distribution of proteins sharing common functional labels were observed for both enzyme/non-enzyme classification and first-level EC categories.

### 3.2 Benchmarking against existing EC enzyme prediction tools

To assess the performance of ESM-ECForest, we compared it with three representative sequence-based EC prediction methods: DeepEC, ProteInfer, and DeepECTransformer. Together, these methods cover convolutional neural network-, dilated convolutional network-, and transformer-based architectures that are widely used for enzyme function prediction.

#### 3.2.1 Overall benchmark performance

We compared ESM-ECForest with DeepECTransformer, ProteInfer, and DeepEC across the four levels of the EC hierarchy (Fig. 5). Full benchmarking results are provided in Supplementary Table S9. ESM-ECForest achieved the highest weighted F1 score at every level, reaching 0.944, 0.929, 0.920, and 0.902 from Levels 1 to 4, respectively (Fig. 5A). In comparison, DeepECTransformer achieved weighted F1 scores of 0.925–0.857, ProteInfer 0.918–0.821, and DeepEC 0.660–0.548 across the same hierarchy.

**Fig. 5.**
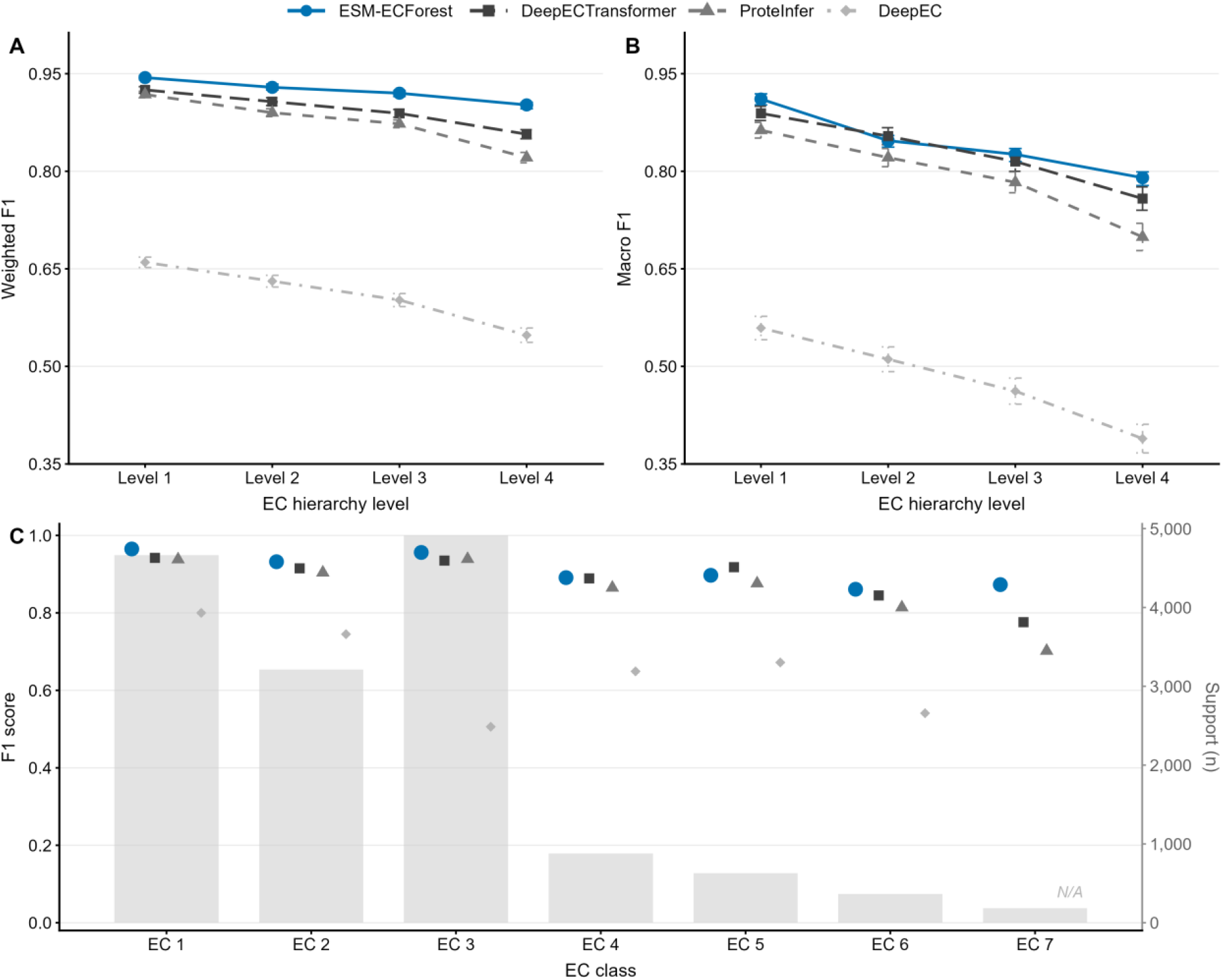
Benchmarking of ESM-ECForest against DeepECTransformer, ProteInfer, and DeepEC. (A) Weighted F1 scores across EC hierarchy Levels 1–4. (B) Macro F1 scores across EC hierarchy Levels 1–4. (C) Class-wise F1 scores at EC Level 1. Grey bars indicate the number of proteins in each EC class (right y-axis). Error bars denote 95% bootstrap confidence intervals. EC class 7 is not supported by the original DeepEC implementation and is shown as N/A. Exact numerical values underlying all panels are provided in Supplementary Table S9.

Performance differences increased with EC hierarchy depth. Between Levels 1 and 4, weighted F1 decreased by 0.042 for ESM-ECForest, compared with declines of 0.068, 0.097, and 0.112 for DeepECTransformer, ProteInfer, and DeepEC, respectively. At Level 4, ESM-ECForest exceeded DeepECTransformer and ProteInfer by 0.045 and 0.081 weighted F1 units, respectively.

A similar trend was observed for macro F1 scores (Fig. 5B). ESM-ECForest achieved macro F1 values of 0.911, 0.847, 0.826, and 0.790 across Levels 1–4. Although DeepECTransformer slightly outperformed ESM-ECForest at Level 2 (0.854 vs. 0.847), ESM-ECForest achieved the highest macro F1 at the remaining levels. At Level 4, its macro F1 remained higher than that of DeepECTransformer (0.758), ProteInfer (0.699), and DeepEC (0.389).

Class-wise evaluation at EC Level 1 further demonstrated consistent performance across enzyme categories (Fig. 5C). ESM-ECForest achieved F1 scores of 0.965, 0.932, and 0.956 for the three most abundant classes (EC 1–3), and 0.861-0.897 for the less represented classes (EC 4–7). Among the seven EC classes, ESM-ECForest achieved the highest F1 score in six, with DeepECTransformer performing slightly better only for EC class 5 (0.918 vs. 0.897). The largest advantage was observed for EC class 7, where ESM-ECForest achieved an F1 score of 0.873 compared with 0.776 for DeepECTransformer and 0.702 for ProteInfer. Predictions for EC class 7 were not available from the original DeepEC implementation.

#### 3.2.2 Runtime and memory comparison

Runtime and peak memory usage were evaluated using the benchmark FASTA dataset. All methods were executed with 24 CPU threads and without GPU acceleration. DeepECTransformer completed prediction in 6.4 min with a peak memory usage of 9.1 GB, followed by DeepEC (8.6 min, 18.9 GB). ESM-ECForest required 11.6 min and 13.4 GB RAM, whereas ProteInfer showed the highest runtime and memory consumption (44.2 min and 36.4 GB, respectively) (Table 1).

**Table 1.**
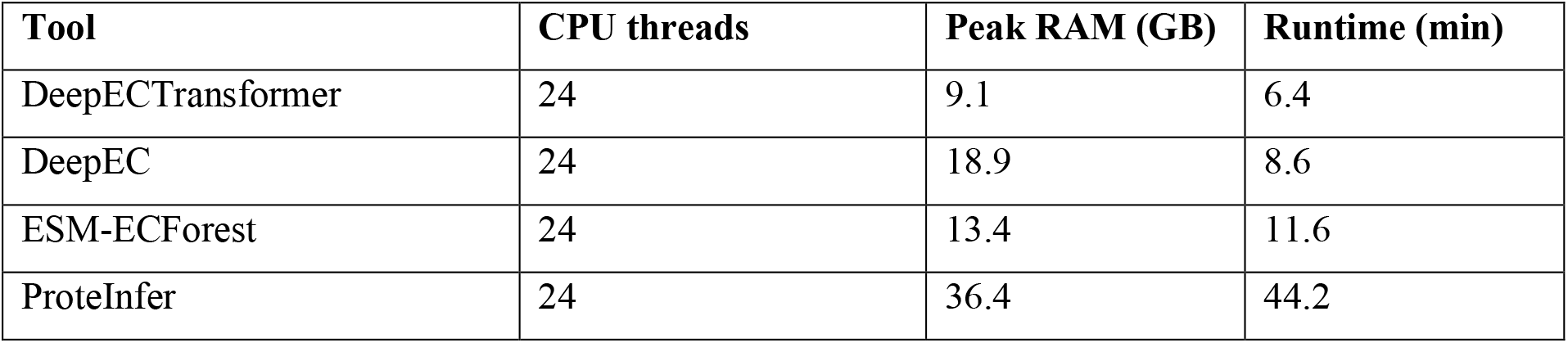
Runtime and memory usage of enzyme annotation tools.

Overall, ESM-ECForest showed moderate computational requirements relative to the other methods evaluated. Although it was slower than DeepECTransformer and DeepEC, its runtime and memory usage remained within a practical range for benchmark-scale enzyme annotation. ProteInfer required substantially greater computational resources, which may partly reflect its default workflow that generates Pfam and Gene Ontology annotations in addition to EC predictions.

Runtime was measured for prediction on the benchmark FASTA dataset. All methods were run using 24 CPU threads without GPU acceleration. Peak RAM indicates the maximum memory usage recorded during prediction. ProteInfer was evaluated using its default workflow, which additionally generates Pfam and Gene Ontology annotations.

## 4 Discussion

The rapid expansion of protein sequence databases continues to outpace the experimental characterization of enzyme functions, creating a persistent gap between sequence availability and functional annotation. Our results indicate that pretrained protein language model embeddings, combined with a two-stage prediction framework, can effectively support hierarchical EC classification. The performance gains observed across multiple EC levels suggest that language-model-derived representations encode biologically meaningful features relevant to enzyme function. Nevertheless, the model does not perform uniformly across all EC categories, and reduced accuracy in certain classes points to intrinsic challenges associated with functional heterogeneity and data scarcity. These findings provide insight into both the strengths and current limitations of embedding-based enzyme annotation approaches and are relevant for applications that depend on accurate enzyme function prediction, including genome-scale metabolic reconstruction.

### 4.1 Protein language model embeddings enable robust hierarchical enzyme prediction

Our results show that strong hierarchical EC prediction performance can be achieved using pretrained sequence representations combined with a relatively lightweight downstream classifier. Although ESM-ECForest uses a relatively simple Random Forest classifier, its performance remained stable across successive EC levels, with only a moderate decrease as the prediction task moved toward more specific functional assignments. The limited performance decline suggests that information relevant to both broad and fine-grained enzyme functions is retained within the ESM-2 embedding space.

This interpretation is also reflected in the UMAP projection. Enzyme and non-enzyme sequences were distributed in partially distinct regions, while enzymes belonging to the same first-level EC category showed a tendency to cluster together. Complete separation would not be expected given the considerable functional and evolutionary overlap among enzyme families. Nevertheless, the observed structure indicates that functionally related proteins occupy similar regions of the embedding space before any task-specific training is performed [32,33,46].

An important component of the framework is the initial discrimination between enzymes and non-enzymes. Stage A achieved an F1 score of 0.976 on the independent benchmark, with similarly high precision and recall, indicating that ESM-2 representations contain sufficient information to distinguish catalytic from non-catalytic proteins even under the homology-aware evaluation used here. This initial filtering step is particularly relevant for end-to-end annotation because false-negative predictions at Stage A cannot subsequently receive an EC assignment, whereas false-positive predictions are passed to Stage B and may generate inappropriate EC annotations. The low error rate observed at Stage A therefore limits the propagation of classification errors into the downstream EC-prediction step and supports the use of an identify-then-classify architecture.

The model showed relatively consistent behavior across different EC classes, including EC 4, EC 6, and EC 7, which are frequently associated with greater sequence heterogeneity and comparatively limited annotation resources [16]. Performance in these categories may reflect the ability of pretrained embeddings to capture properties that are not restricted to individual enzyme families. Instead of depending primarily on family-specific sequence patterns, the representation appears to preserve broader functional signals that remain informative across diverse enzyme groups [29].

This observation is unlikely to be specific to the present model and may instead reflect properties of protein language model representations that have been reported previously[38]. Representations generated by ESM-2 have been associated with structural, evolutionary, and functional characteristics of proteins, despite being trained without explicit functional supervision [32]. The success of ESMFold, in line with earlier studies on unsupervised protein language modeling [33,47,48], further illustrated the extent to which information embedded in protein language model representations can learn representations reflecting biological structure and function and support downstream biological inference. In the context of enzyme annotation, the present results suggest that these embeddings contain sequence-level constraints related to catalytic function, domain architecture, and protein structure, providing a useful basis for hierarchical EC prediction [34].

### 4.2 Why ESM embeddings help

A notable characteristic of ESM-ECForest is its separation of protein representation learning from supervised enzyme classification. Rather than deriving sequence features directly from EC annotations, the framework first generates protein embeddings using the pretrained ESM-2 encoder and then applies Random Forest classifiers to the resulting representations. In this way, learning general properties of protein sequences and learning function-specific decision boundaries are treated as two related but distinct tasks.

One possible explanation for the effectiveness of ESM-derived representations lies in the pretraining objective. ESM-2 is trained to recover masked amino acids from their surrounding sequence context:

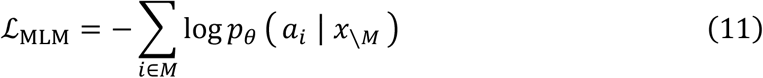

where ℒ_MLM_ is the masked language modelling loss used to train ESM-2, *x* = (*a*_1_, *a*_2_, …, *a*_*L*_) denotes an amino acid sequence of length *L, M* denotes the set of masked positions, *a*_*i*_ is the true amino-acid at position *i*, and *p*_*θ*_(*a*_*i*_ ∣ *x*_\*M*_) is the probability that the model assigns to the correct amino acid *a*_*i*_ given the surrounding sequence. Optimizing this objective requires the model to integrate information across multiple sequence positions rather than relying only on local patterns.As a result, the learned representations can capture constraints associated with protein structure, evolutionary conservation, and biochemical function [33,34]. Self-attention may further allow information from distant positions in the primary sequence to contribute to the representation, which is relevant because catalytic residues can be separated in sequence while becoming spatially adjacent in the folded protein.

In ESM-ECForest, the final-layer BOS token representation was used as a fixed-length sequence-level embedding, with chunk-level embeddings averaged only for sequences longer than the ESM-2 input limit. Random Forest classifiers were then trained on these embeddings to distinguish enzymes from non-enzymes and to assign EC labels. Conceptually, the two-stage prediction procedure can be written as:

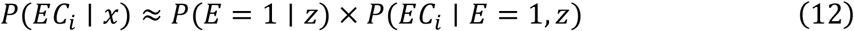

where *E* = 1 denotes the event that a protein is enzymatic, *z* denotes the fixed-length ESM-2 sequence embedding. The first stage estimates whether a sequence is likely to encode an enzyme, whereas the second stage assigns EC numbers only to sequences classified as enzymatic. This two-stage framework separates enzyme identification from EC classification and limits EC assignment to proteins predicted as enzymes.

### 4.3 Benchmarking reveals improved robustness over existing sequence-based tools

The benchmarking results highlight several differences among DeepEC, ProteInfer, DeepECTransformer, and ESM-ECForest, which employ distinct strategies for extracting functional information from protein sequences. DeepEC is primarily based on convolutional neural networks and supplements prediction with homology analysis when confidence is insufficient [20]. ProteInfer adopts dilated convolutions for alignment-free EC prediction [24], whereas DeepECTransformer performs supervised EC classification using transformer architectures [21]. These approaches differ mainly in how sequence context is represented and incorporated into prediction.

For a standard one-dimensional convolution, the output of a hidden unit can be expressed as:

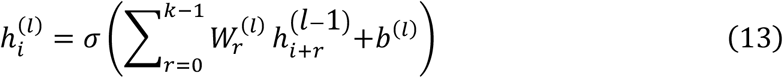

where *k* is the kernel size, 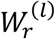 is the convolutional weight at position *r*, and 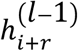 represents a neighbouring input feature, *b*^(*l*)^ is the bias term, and *σ* is the activation function. Convolutional operations are effective at detecting local sequence patterns, including motifs associated with catalytic residues, cofactor binding sites, and enzyme-family signatures [18,38]. However, information is aggregated within a finite receptive field, making long-range sequence relationships more difficult to capture without increasing network depth. ProteInfer addresses this limitation through dilated convolutions, which expand the receptive field without increasing the kernel size [49]. The enlarged receptive field enables information to be incorporated from more distant sequence positions while maintaining efficient inference. Nevertheless, interactions are still determined by predefined convolutional pathways. Consequently, dependencies between residues are inferred indirectly through successive local operations.

Transformer-based models adopt a different mechanism. In self-attention, the contribution of residue *j* to the representation of residue *i* is determined dynamically by the input sequence:

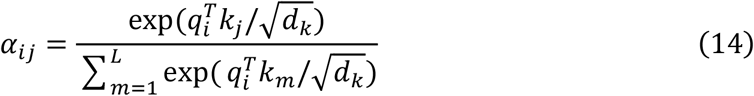

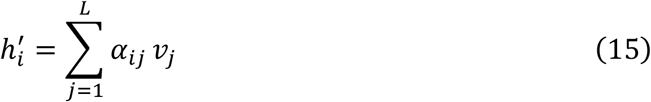

where *α*_*ij*_ is the attention weight describing how strongly residue *i* attends to residue *j*, 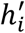 is the updated context-dependent representation of residue *i* after the attention operation, and *q*_*i*_, *k*_*j*_, and *v*_*j*_ denote the query, key, and value representations, respectively. Unlike convolutional operations, the attention weights are not fixed after training but vary according to sequence context. This property allows transformer models to model relationships among residues regardless of their sequence separation. Such interactions are particularly relevant for enzymes, whose catalytic properties often depend on spatially coordinated residues distributed across different regions of the primary sequence.

DeepECTransformer may benefits from this attention-based formulation and is therefore expected to capture long-range functional signals more effectively than conventional CNN architectures[21,50]. However, as an end-to-end supervised model, its performance remains closely linked to the quantity and distribution of available EC annotations. This dependency can become limiting when training examples are sparse or unevenly distributed across functional classes.

ESM-ECForest differs from the above approaches in that sequence representation learning is performed prior to EC classification. The transformer encoder is trained on large-scale protein sequence corpora before being applied to the enzyme prediction task. As a result, the downstream classifier operates on embeddings that already contain information learned from evolutionary and structural patterns present in protein sequences. In contrast, CNN-based approaches and supervised transformers must learn a larger fraction of the sequence-to-function mapping directly from labelled training data.

This distinction may partly account for the behaviour observed in the benchmark experiments. The performance advantage of ESM-ECForest was most apparent in EC classes such as EC 4, EC 6, and EC 7, which are characterised by substantial functional diversity and comparatively limited annotation resources [4,16,34]. Under these conditions, local motif detection alone may not always provide sufficient discriminatory information. Pretrained embeddings may therefore offer an advantage by positioning proteins with related structural or evolutionary constraints closer together in representation space, even when sequence similarity is relatively low [32,33]. The downstream Random Forest classifier can subsequently identify non-linear class boundaries within this representation space using comparatively few task-specific parameters.

Runtime differed among the evaluated tools. DeepECTransformer and DeepEC completed predictions faster than ESM-ECForest, whereas ProteInfer required the longest runtime and highest memory usage. The additional runtime of ESM-ECForest was primarily associated with ESM-2 embedding generation. Despite this overhead, ESM-ECForest achieved the highest prediction performance across all EC levels.

It should also be noted that differences among tools are influenced by factors beyond model architecture alone. Variations in training datasets, EC coverage, label curation procedures, confidence thresholds, and auxiliary prediction modules can all affect benchmark outcomes. For example, DeepEC does not provide predictions for EC 7 enzymes, which inevitably affects comparative performance when translocases are included in the evaluation. The observed differences should therefore not be interpreted solely as evidence for the superiority of one modelling strategy over another. Instead, the benchmark results suggest that combining pretrained protein embeddings with a lightweight supervised classifier provides an effective framework for hierarchical EC annotation under the datasets and evaluation settings considered in this study.

### 4.4 Biochemical complexity limits prediction in EC 6 and EC 7

Despite the overall improvement achieved by ESM-ECForest, EC 6 and EC 7 remained among the most difficult categories to predict accurately. This trend is unlikely to arise solely from limitations of the prediction model and may instead reflect intrinsic differences in the biological characteristics of these enzyme classes. Ligases (EC 6) catalyse bond-forming reactions that are typically coupled to energy consumption, whereas translocases (EC 7) mediate the movement of ions or molecules across membranes or their separation within membrane systems. In addition to the diversity of transported substrates, EC 7 enzymes are further subdivided according to the energetic mechanisms that drive transport [51].

A major challenge is the substantial functional heterogeneity present within both classes. Enzymes assigned to the same top-level EC category may differ considerably in substrate specificity, catalytic mechanism, cofactor requirements, and domain organization. Consequently, proteins sharing an EC 6 or EC 7 designation do not necessarily exhibit strong sequence similarity, making sequence-based discrimination inherently more difficult than for enzyme groups with more conserved functional architectures.

The challenge is particularly apparent for translocases. Many EC 7 enzymes are membrane-associated proteins or components of multi-subunit complexes whose activities depend on factors extending beyond the primary amino acid sequence. Transmembrane topology, oligomeric state, cellular localization, electrochemical gradients, and coupling to ATP hydrolysis or redox processes can all contribute to biological function. Although some of this information is reflected indirectly in sequence patterns, it may not be fully captured by sequence embeddings alone, especially when different transport systems employ distinct structural solutions to achieve similar functions [34,52].

Dataset characteristics may further contribute to the observed performance differences. Compared with oxidoreductases, transferases, or hydrolases, EC 6 and EC 7 enzymes are represented by fewer experimentally characterized examples in many public annotation resources [16,53]. EC 6 and EC 7 were represented by 995 proteins in our training dataset, compared with 13,667 for the more abundant EC classes. Reduced annotation density limits the diversity of functional patterns available during model training and increases the influence of incomplete or uncertain reference labels. Under such conditions, benchmark performance may reflect not only prediction errors but also limitations of the available annotations themselves.

Taken together, the residual errors observed for EC 6 and EC 7 are consistent with the underlying biochemical complexity of these enzyme classes. Further improvements will likely require the integration of information beyond primary sequence, including structural features, membrane topology, active-site and ligand-binding information, domain composition, protein-complex organization, and reaction-level biochemical constraints [34]. Such information may help resolve functional distinctions that are difficult to infer from sequence data alone.

### 4.5 Implications for metabolic annotation and genome-scale modeling

The main practical value of EC prediction is that it links protein sequences to biochemical reactions. EC numbers provide a functional bridge from gene products to reaction-level annotations, which form the basis of gene-protein-reaction associations in genome-scale metabolic models (GEMs) [54–56]. GEMs represent organism-wide metabolic networks and can be used to simulate metabolic fluxes under different genetic or environmental conditions. By connecting genes to metabolic reactions, these annotations enable the systematic reconstruction and analysis of cellular metabolic networks.

In microbial systems, improved enzyme annotation can facilitate several stages of GEM development [55]. One immediate application is the functional characterization of proteins that remain incompletely annotated. Predicted EC assignments may also assist in identifying candidate genes associated with missing metabolic reactions during gap-filling procedures [57]. In addition, improved enzyme annotations can support the refinement of existing GPR relationships by providing evidence for alternative or previously unrecognized enzyme candidates. Such applications are particularly relevant for *Saccharomyces cerevisiae*, where community-curated reconstructions such as Yeast9 and yeast-GEM continue to serve as important resources for metabolic engineering, and multi-omics integration [55,58,59].

At the same time, the connection between EC annotation and cellular metabolism should not be overstated. An EC prediction reflects the potential catalytic activity encoded by a protein sequence rather than its realised biological function under a specific condition. Whether an enzyme contributes to metabolic flux in vivo depends on additional factors, including gene expression, protein abundance, subcellular localization, substrate availability, regulatory mechanisms, and environmental context. Consequently, EC predictions are most informative when considered alongside complementary sources of biological evidence.

From this perspective, ESM-ECForest is perhaps best viewed as a prioritization tool rather than a definitive annotation system. The predicted EC assignments can be used to guide subsequent analyses and to generate hypotheses regarding enzyme function, particularly for poorly characterized proteins. Integration with transcriptomic, proteomic, metabolomic, phenotypic, and curated biochemical databases will remain necessary before such predictions can be incorporated into high-confidence metabolic reconstructions. Nevertheless, improvements in sequence-based enzyme annotation may help narrow the gap between the rapidly growing number of available protein sequences and the slower accumulation of experimentally validated metabolic knowledge.

### 4.6 Limitations and future directions

One limitation of the current framework arises from the EC system itself. Although EC numbers provide a standardized description of enzyme activity, they do not always capture the full functional diversity of enzymes [3,4]. Proteins sharing the same EC assignment may differ substantially in substrate preference, cofactor dependence, reaction conditions, cellular localization, or physiological role. Conversely, a single EC designation may encompass multiple closely related biochemical reactions. As a result, accurate EC prediction does not necessarily translate into reaction-level resolution, which is often required for metabolic reconstruction and functional interpretation.

This limitation highlights a broader challenge in enzyme annotation. Future developments are likely to require representations that extend beyond EC classification alone and incorporate additional layers of biochemical information [60]. Integrating substrate specificity, cofactor requirements, pathway context, cellular compartmentation, and reaction-level resources such as Rhea could help establish a more direct link between protein sequences and biochemical reactions [56]. Such efforts are consistent with recent attempts to move beyond traditional EC-based enzyme annotation and to represent enzyme function in a more flexible and biologically informative manner [16,34,38,60]. One example is the ECxit framework, which aims to redesign enzyme function annotation through a hierarchical deep-learning-based approach [61].

Several limitations of the present evaluation should also be considered. ESM-ECForest was trained using selected prokaryotic and unicellular eukaryotic proteins from UniProtKB/Swiss-Prot, and its performance may not generalize equally well to other taxonomic groups or poorly represented enzyme families. In addition, the individual contributions of the ESM-2 embeddings, Random Forest classifiers, and two-stage architecture were not evaluated through ablation analyses. Because Stage B classifiers are restricted to EC labels observed during training, ESM-ECForest cannot assign previously unseen terminal EC numbers. However, higher-level EC annotations may still be recovered when predicted and reference EC numbers share the same higher-level classifications. Future evaluations should also examine performance as a function of sequence identity to the training set and assess the ability of the model to recover partial or complete EC annotations that were absent from the training data.

There is also scope for methodological improvements. Although the current framework performs well across multiple EC levels, the confidence associated with individual predictions remains difficult to interpret. Improved probability calibration may be particularly valuable in applications where predictions are used to prioritize experimental validation or support manual curation. In addition, the persistent difficulty of classes such as EC 6 and EC 7 suggests that sequence information alone may not always be sufficient. Structural models, active-site annotations, transmembrane topology, domain organization, and protein-complex information could provide complementary evidence for distinguishing functionally similar enzymes [34].

More broadly, the results underscore the value of pretrained protein representations for large-scale enzyme annotation. At the same time, the remaining prediction errors indicate that enzyme function cannot always be inferred from sequence alone. A natural next step will be to combine sequence-derived representations with structural, biochemical, and systems-level information, thereby moving from EC prediction toward functionally resolved and model-ready metabolic annotations.

## Acknowledgements

We thank the division of Biotechnology and Applied Microbiology at Lund University and especially the Metabolic Engineering group for their interesting comments on the work.

## Funding

This work was supported by Lund University and the Faculty of Engineering at Lund University, as well as by the Sten K Johnsons Foundation.

## Author contributions

X.H. and G.G. conceptualized the work. X.H. performed the analysis and wrote the manuscript. G.G. contributed to the writing and revision of the manuscript.

## Competing interests

The authors declare that they have no competing interests.

## Data Availability

All data supporting the findings of this study are available within the manuscript and its Supplementary Materials. The source code required to reproduce the analyses is available at GitHub (https://github.com/Xiao88866/ESM-ECForest). The trained models generated during this study are publicly available through Zenodo (https://doi.org/10.5281/zenodo.21699471).

## Notes

### Competing Interest Statement

The authors have declared no competing interest.

## References

[1] Morgat A, Lombardot T, Coudert E, Axelsen K, Neto TB, Gehant S, et al. Enzyme annotation in UniProtKB using Rhea. Bioinformatics 2020;36:1896–901. 10.1093/bioinformatics/btz817.

[2] Kanehisa M. Enzyme Annotation and Metabolic Reconstruction Using KEGG. In: Kihara D, editor. Protein Function Prediction: Methods and Protocols, New York, NY: Springer; 2017, p. 135–45. 10.1007/978-1-4939-7015-5_11.

[3] McDonald AG, Tipton KF. Fifty-five years of enzyme classification: advances and difficulties. The FEBS Journal 2014;281:583–92. 10.1111/febs.12530.

[4] McDonald AG, Tipton KF. Enzyme nomenclature and classification: the state of the art. The FEBS Journal 2023;290:2214–31. 10.1111/febs.16274.

[5] Cornish-Bowden A. Current IUBMB recommendations on enzyme nomenclature and kinetics. Perspectives in Science 2014;1:74–87. 10.1016/j.pisc.2014.02.006.

[6] Qian W, Wang X, Kang Y, Pan P, Hou T, Hsieh C-Y. A general model for predicting enzyme functions based on enzymatic reactions. J Cheminform 2024;16:38. 10.1186/s13321-024-00827-y.

[7] Rong D, Zhong B, Zheng W, Hong L, Liu N. Autoregressive enzyme function prediction with multi-scale multi-modality fusion. Brief Bioinform 2025;26:bbaf476. 10.1093/bib/bbaf476.

[8] Quester S, Schomburg D. EnzymeDetector: an integrated enzyme function prediction tool and database. BMC Bioinformatics 2011;12:376. 10.1186/1471-2105-12-376.

[9] Radivojac P, Clark WT, Oron TR, Schnoes AM, Wittkop T, Sokolov A, et al. A large-scale evaluation of computational protein function prediction. Nat Methods 2013;10:221–7. 10.1038/nmeth.2340.

[10] Altschul SF, Gish W, Miller W, Myers EW, Lipman DJ. Basic Local Alignment Search Tool. J Mol Biol 1990;215:403–10. 10.1016/S0022-2836(05)80360-2.

[11] Hu G, Kurgan L. Sequence Similarity Searching. Current Protocols in Protein Science 2019;95:e71. 10.1002/cpps.71.

[12] Krogh A, Brown M, Mian IS, Sjölander K, Haussler D. Hidden Markov models in computational biology. Applications to protein modeling. J Mol Biol 1994;235:1501–31. 10.1006/jmbi.1994.1104.

[13] Mor B, Garhwal S, Kumar A. A Systematic Review of Hidden Markov Models and Their Applications. Arch Computat Methods Eng 2021;28:1429–48. 10.1007/s11831-020-09422-4.

[14] Holm L, Sander C. Protein Structure Comparison by Alignment of Distance Matrices. J Mol Biol 1993;233:123–38. 10.1006/jmbi.1993.1489.

[15] Shindyalov IN, Bourne PE. A database and tools for 3-D protein structure comparison and alignment using the Combinatorial Extension (CE) algorithm. Nucleic Acids Res 2001;29:228–9. 10.1093/nar/29.1.228.

[16] Yu T, Cui H, Li JC, Luo Y, Jiang G, Zhao H. Enzyme function prediction using contrastive learning. Science 2023;379:1358–63. 10.1126/science.adf2465.

[17] Yu C, Zavaljevski N, Desai V, Reifman J. Genome-wide enzyme annotation with precision control: Catalytic families (CatFam) databases. Proteins 2009;74:449–60. 10.1002/prot.22167.

[18] Li Y, Wang S, Umarov R, Xie B, Fan M, Li L, et al. DEEPre: sequence-based enzyme EC number prediction by deep learning. Bioinformatics 2018;34:760–9. 10.1093/bioinformatics/btx680.

[19] Nursimulu N, Xu LL, Wasmuth JD, Krukov I, Parkinson J. Improved enzyme annotation with EC-specific cutoffs using DETECT v2. Bioinformatics 2018;34:3393–5. 10.1093/bioinformatics/bty368.

[20] Ryu JY, Kim HU, Lee SY. Deep learning enables high-quality and high-throughput prediction of enzyme commission numbers. Proc Natl Acad Sci USA 2019;116:13996–4001. 10.1073/pnas.1821905116.

[21] Kim GB, Kim JY, Lee JA, Norsigian CJ, Palsson BO, Lee SY. Functional annotation of enzyme-encoding genes using deep learning with transformer layers. Nat Commun 2023;14:7370. 10.1038/s41467-023-43216-z.

[22] Dalkiran A, Rifaioglu AS, Martin MJ, Cetin-Atalay R, Atalay V, Doğan T. ECPred: a tool for the prediction of the enzymatic functions of protein sequences based on the EC nomenclature. BMC Bioinformatics 2018;19:334. 10.1186/s12859-018-2368-y.

[23] Kumar N, Skolnick J. EFICAz2.5: application of a high-precision enzyme function predictor to 396 proteomes. Bioinformatics 2012;28:2687–8. 10.1093/bioinformatics/bts510.

[24] Sanderson T, Bileschi ML, Belanger D, Colwell LJ. ProteInfer, deep neural networks for protein functional inference. eLife 2023;12:e80942. 10.7554/eLife.80942.

[25] Claudel-Renard C, Chevalet C, Faraut T, Kahn D. Enzyme-specific profiles for genome annotation: PRIAM. Nucleic Acids Res 2003;31:6633–9. 10.1093/nar/gkg847.

[26] Busi R, Machingal P, Hemachandra N, Balaji PV. Systematic assessment of homology-based methods for fine-grained functional annotation using diverse protein families 2025:2025.09.17.676781. 10.1101/2025.09.17.676781.

[27] Kaminski K, Ludwiczak J, Pawlicki K, Alva V, Dunin-Horkawicz S. pLM-BLAST: distant homology detection based on direct comparison of sequence representations from protein language models. Bioinformatics 2023;39:btad579. 10.1093/bioinformatics/btad579.

[28] Rost B. Twilight zone of protein sequence alignments. Protein Eng 1999;12:85–94. 10.1093/protein/12.2.85.

[29] Capela J, Zimmermann-Kogadeeva M, van Dijk ADJ, de Ridder D, Dias O, Rocha M. Comparative Assessment of Protein Large Language Models for Enzyme Commission Number Prediction. BMC Bioinformatics 2025;26:68. 10.1186/s12859-025-06081-9.

[30] Vieira LC, Handojo ML, Wilke CO. Medium-sized protein language models perform well at transfer learning on realistic datasets. Sci Rep 2025;15:21400. 10.1038/s41598-025-05674-x.

[31] Xiao Y, Zhao W, Zhang J, Jin Y, Zhang H, Ren Z, et al. Protein Large Language Models: A Comprehensive Survey. In: Christodoulopoulos C, Chakraborty T, Rose C, Peng V, editors. Findings of the Association for Computational Linguistics: EMNLP 2025, Suzhou, China: Association for Computational Linguistics; 2025, p. 23080–103. 10.18653/v1/2025.findings-emnlp.1255.

[32] Lin Z, Akin H, Rao R, Hie B, Zhu Z, Lu W, et al. Evolutionary-scale prediction of atomic-level protein structure with a language model. Science 2023;379:1123–30. 10.1126/science.ade2574.

[33] Rives A, Meier J, Sercu T, Goyal S, Lin Z, Liu J, et al. Biological structure and function emerge from scaling unsupervised learning to 250 million protein sequences. Proc Natl Acad Sci USA 2021;118:e2016239118. 10.1073/pnas.2016239118.

[34] Song Y, Yuan Q, Chen S, Zeng Y, Zhao H, Yang Y. Accurately predicting enzyme functions through geometric graph learning on ESMFold-predicted structures. Nat Commun 2024;15:8180. 10.1038/s41467-024-52533-w.

[35] Sathyamoorthy R, Puri M. Protein Language Models Outperform BLAST for Evolutionarily Distant Enzymes: A Systematic Benchmark of EC Number Prediction 2026:2026.03.31.715487. 10.64898/2026.03.31.715487.

[36] Fan W, Zhou Y, Wang S, Yan Y, Liu H, Zhao Q, et al. Computational Protein Science in the Era of Large Language Models (LLMs) 2025. 10.48550/arXiv.2501.10282.

[37] Chen J-Y, Wang J-F, Hu Y, Li X-H, Qian Y-R, Song C-L. Evaluating the advancements in protein language models for encoding strategies in protein function prediction: a comprehensive review. Front Bioeng Biotechnol 2025;13:1506508. 10.3389/fbioe.2025.1506508.

[38] Tan Q, Xiao J, Chen J, Wang Y, Zhang Z, Zhao T, et al. ifDEEPre: large protein language-based deep learning enables interpretable and fast predictions of enzyme commission numbers. Brief Bioinform 2024;25:bbae225. 10.1093/bib/bbae225.

[39] The UniProt Consortium. UniProt: a hub for protein information. Nucleic Acids Res 2015;43:D204–12. 10.1093/nar/gku989.

[40] Breiman L. Random Forests. Machine Learning 2001;45:5–32. 10.1023/A:1010933404324.

[41] Díaz-Uriarte R, Alvarez de Andrés S. Gene selection and classification of microarray data using random forest. BMC Bioinformatics 2006;7:3. 10.1186/1471-2105-7-3.

[42] Chen X, Ishwaran H. Random forests for genomic data analysis. Genomics 2012;99:323–9. 10.1016/j.ygeno.2012.04.003.

[43] Sokolova M, Lapalme G. A systematic analysis of performance measures for classification tasks. Information Processing & Management 2009;45:427–37. 10.1016/j.ipm.2009.03.002.

[44] Tsoumakas G, Katakis I. Multi-Label Classification: An Overview. IJDWM 2007;3:1–13. 10.4018/jdwm.2007070101.

[45] McInnes L, Healy J, Melville J. UMAP: Uniform Manifold Approximation and Projection for Dimension Reduction 2018. 10.48550/arXiv.1802.03426.

[46] Becht E, McInnes L, Healy J, Dutertre C-A, Kwok IWH, Ng LG, et al. Dimensionality reduction for visualizing single-cell data using UMAP. Nat Biotechnol 2019;37:38–44. 10.1038/nbt.4314.

[47] Alley EC, Khimulya G, Biswas S, AlQuraishi M, Church GM. Unified rational protein engineering with sequence-based deep representation learning. Nat Methods 2019;16:1315–22. 10.1038/s41592-019-0598-1.

[48] Rao R, Bhattacharya N, Thomas N, Duan Y, Chen X, Canny J, et al. Evaluating Protein Transfer Learning with TAPE. Adv Neural Inf Process Syst 2019;32:9689–701.

[49] Yu F, Koltun V. Multi-Scale Context Aggregation by Dilated Convolutions 2016. 10.48550/arXiv.1511.07122.

[50] Vaswani A, Shazeer N, Parmar N, Uszkoreit J, Jones L, Gomez AN, et al. Attention is All you Need. Adv Neural Inf Process Syst 2017;30:5998–6008.

[51] Saier MH, Reddy VS, Tamang DG, Västermark A. The transporter classification database. Nucleic Acids Res 2014;42:D251–8. 10.1093/nar/gkt1097.

[52] Yang Y, Jerger A, Feng S, Wang Z, Brasfield C, Cheung MS, et al. Improved enzyme functional annotation prediction using contrastive learning with structural inference. Commun Biol 2024;7:1690. 10.1038/s42003-024-07359-z.

[53] Davoudi S, Henry CS, Miller CS, Banaei-Kashani F. EC-Bench: a benchmark for enzyme commission number prediction. Bioinformatics Advances 2026;6:vbag004. 10.1093/bioadv/vbag004.

[54] Orth JD, Thiele I, Palsson BØ. What is flux balance analysis? Nat Biotechnol 2010;28:245–8. 10.1038/nbt.1614.

[55] Gu C, Kim GB, Kim WJ, Kim HU, Lee SY. Current status and applications of genome-scale metabolic models. Genome Biol 2019;20:121. 10.1186/s13059-019-1730-3.

[56] Bansal P, Morgat A, Axelsen KB, Muthukrishnan V, Coudert E, Aimo L, et al. Rhea, the reaction knowledgebase in 2022. Nucleic Acids Res 2022;50:D693–700. 10.1093/nar/gkab1016.

[57] Thiele I, Palsson BØ. A protocol for generating a high-quality genome-scale metabolic reconstruction. Nat Protoc 2010;5:93–121. 10.1038/nprot.2009.203.

[58] Nookaew I, Olivares-Hernández R, Bhumiratana S, Nielsen J. Genome-scale metabolic models of Saccharomyces cerevisiae. Methods Mol Biol 2011;759:445–63. 10.1007/978-1-61779-173-4_25.

[59] Lu H, Li F, Sánchez BJ, Zhu Z, Li G, Domenzain I, et al. A consensus S. cerevisiae metabolic model Yeast8 and its ecosystem for comprehensively probing cellular metabolism. Nat Commun 2019;10:3586. 10.1038/s41467-019-11581-3.

[60] Shi Z, Zhu J, Wang D, Chen B, Yuan Q, Mao Z, et al. RXNRECer Enables Fine-grained Enzymatic Function Annotation through Active Learning and Protein Language Models 2026. 10.48550/arXiv.2603.12694.

[61] ECxit. Inria n.d. https://www.inria.fr/en/ecxit (accessed July 23, 2026).

