## Supplementary Information for "A Two-Stage ESM-Based Machine Learning Pipeline for Robust Hierarchical Enzyme Function Prediction"

#### Supporting Figures

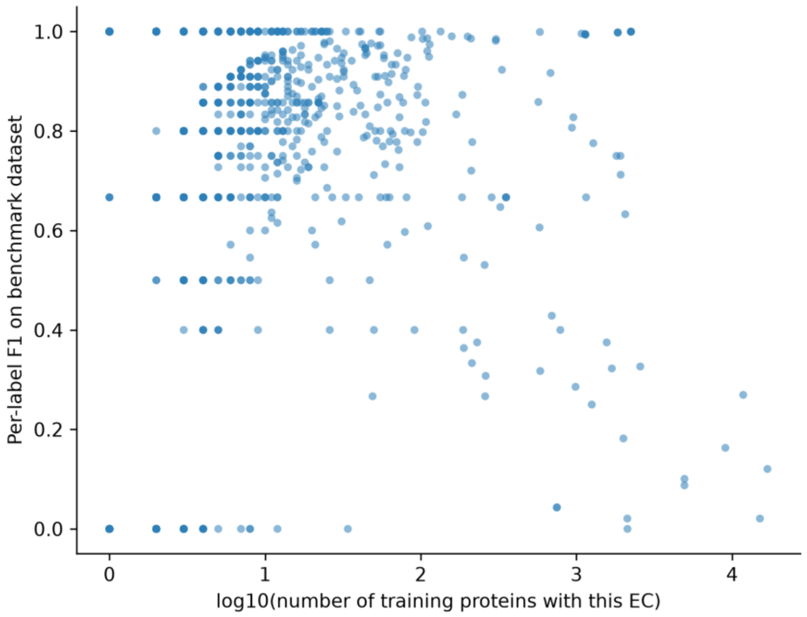


Figure S1: Relationship between EC-label training support and benchmark F1 score. Each point is one EC label. The x-axis is the log10-transformed number of training proteins annotated with that EC; the y-axis is the corresponding per-EC F1 on the benchmark dataset.

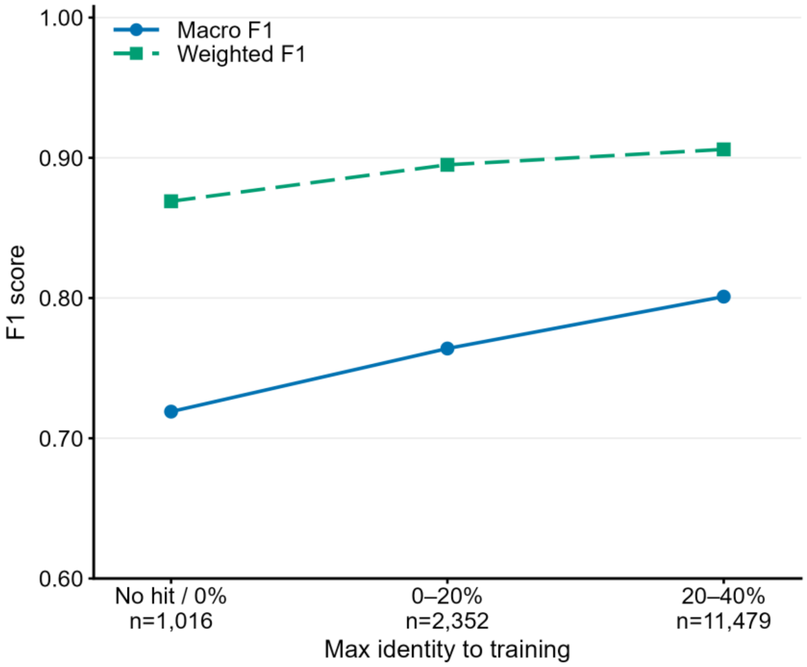


Figure S2: EC Level 4 performance across sequence-identity bins. Proteins were grouped according to their maximum sequence identity to the training dataset. Macro- and weighted-average F1 scores were calculated for each bin.

### Supporting Tables

**Table S1:** Taxonomic groups included in dataset construction.

| Organism group | Taxon ID |
|---|---|
| Bacteria | 2 |
| Archaea | 2157 |
| Saccharomyces | 4930 |
| Schizosaccharomyces | 4895 |
| Tetrahymena | 5890 |
| Trypanosoma | 5690 |
| Leishmania | 5658 |
| Plasmodium | 5820 |
| Giardia | 5740 |
| Entamoeba | 5758 |
| Chlamydomonas | 3052 |
| Euglena | 3038 |
| Dunaliella | 3044 |
| Candida | 5475 |
| Cryptococcus | 5206 |

**Table S2A:** Distribution of enzyme and non-enzyme sequences in the training, validation, and test sets.

| Class | Train | Val | Test | Total |
|---|---|---|---|---|
| Enzyme | 90,704 | 4,615 | 9,329 | 104,648 |
| Non-enzyme | 64,841 | 6,095 | 9,430 | 80,366 |

**Table S2B:** Distribution of EC level 1 annotations in the training, validation, and test sets.

| EC class | Train | Val | Test | Total |
|---|---|---|---|---|
| 1 | 29,572 | 955 | 503 | 31,030 |
| 2 | 34,975 | 10,196 | 1,439 | 46,610 |
| 3 | 16,407 | 2,963 | 486 | 19,856 |
| 4 | 14,087 | 939 | 919 | 15,945 |
| 5 | 2,891 | 112 | 80 | 3,083 |
| 6 | 6,488 | 70 | 97 | 6,655 |
| 7 | 503 | 16 | 24 | 543 |

**Table S3:** Random Forest hyperparameters used in ESM-ECForest.

| Hyperparameter | Stage A enzyme classifier | Stage B EC classifier |
|---|---|---|

| | | |
|---|---|---|
| n_estimators | 400 | 400 for each binary classifier |
| max_depth | 40 | 40 |
| bootstrap | True | True |
| max_samples | 0.8 | 0.8 |
| min_samples_split | 3 | 3 |
| min_samples_leaf | 1 | 1 |
| max_features | ’sqrt’ | ‘sqrt’ |
| random_state | 42 | 42 |
| n_jobs | -1 | -1 |

**Table S4:** Species included in the benchmarking dataset.

| Species | Taxon ID |
|---|---|
| Yarrowia lipolytica | 4952 |
| Komagataella pastoris | 4922 |
| Debaryomyces hansenii | 4959 |
| Dictyostelium discoideum | 44689 |
| Acanthamoeba castellanii | 5755 |
| Toxoplasma gondii | 5811 |
| Paramecium tetraurelia | 5888 |
| Naegleria gruberi | 5762 |
| Micromonas pusilla | 38833 |
| Corynebacterium glutamicum | 1718 |
| Deinococcus radiodurans | 243230 |
| Shewanella oneidensis | 211586 |
| Methanosarcina mazei | 192952 |
| Nitrosopumilus maritimus | 436308 |
| Thermococcus kodakarensis | 69014 |

**Table S5:** Software implementations used for benchmarking.

| Tool | Implementation | Commit |
|---|---|---|
| DeepEC | Bitbucket | adf79c9571d52218302c5323835f3508d88361aa |
| ProteInfer | GitHub | 0acc7b16eb93f21d88c15dcfe3c7828aef2881c8 |
| DeepECTransformer | GitHub | 0eff1da154a31ad2b20909bd4bb48fbf6d55c569 |

**Table S6:** Hardware and software specifications of the benchmarking environment.

| Component | Specification |
|---|---|
| CPU | AMD EPYC 7413 24-Core Processor |
| CPU threads used | 24 |

| | |
|---|---|
| Memory allocated | 200 GB |
| GPU | Not used |
| Python | 3.9.23 |
| PyTorch | 2.7.1+cu126 |

**Table S7:** Performance metrics of Stage A binary enzyme identification

| Metric | Estimate | 95% CI |
|---|---|---|
| Precision | 0.975 | [0.972, 0.977] |
| Recall | 0.977 | [0.975, 0.980] |
| F1 | 0.976 | [0.974, 0.978] |

**Table S8A:** Hierarchical EC prediction performance of ESM-ECForest across EC Levels 1–4.

| Metric | EC level | Macro (95% CI) | Micro (95% CI) | Weighted (95% CI) |
|---|---|---|---|---|
| F1 | 1 | 0.911 [0.902, 0.919] | 0.943 [0.940, 0.946] | 0.944 [0.941, 0.947] |
| F1 | 2 | 0.847 [0.837, 0.855] | 0.921 [0.917, 0.924] | 0.929 [0.925, 0.934] |
| F1 | 3 | 0.826 [0.815, 0.835] | 0.912 [0.908, 0.916] | 0.920 [0.915, 0.924] |
| F1 | 4 | 0.790 [0.778, 0.799] | 0.888 [0.883, 0.892] | 0.902 [0.896, 0.906] |
| Precision | 1 | 0.896 [0.886, 0.904] | 0.952 [0.948, 0.956] | 0.953 [0.949, 0.957] |
| Precision | 2 | 0.864 [0.853, 0.874] | 0.924 [0.918, 0.929] | 0.943 [0.938, 0.947] |
| Precision | 3 | 0.848 [0.837, 0.859] | 0.911 [0.904, 0.917] | 0.932 [0.926, 0.937] |
| Precision | 4 | 0.808 [0.796, 0.819] | 0.882 [0.875, 0.889] | 0.910 [0.903, 0.915] |
| Recall | 1 | 0.928 [0.919, 0.935] | 0.935 [0.932, 0.938] | 0.935 [0.932, 0.938] |
| Recall | 2 | 0.842 [0.832, 0.851] | 0.918 [0.914, 0.922] | 0.918 [0.914, 0.922] |
| Recall | 3 | 0.821 [0.811, 0.832] | 0.913 [0.909, 0.917] | 0.913 [0.909, 0.917] |
| Recall | 4 | 0.785 [0.773, 0.796] | 0.895 [0.890, 0.899] | 0.895 [0.891, 0.899] |

**Table S8B:** Class-specific EC prediction performance of ESM-ECForest at EC Level 1.

| EC class | Precision | Recall | F1 score | Support |
|---|---|---|---|---|
| 1 | 0.983 | 0.948 | 0.965 | 4663 |
| 2 | 0.944 | 0.920 | 0.932 | 3212 |
| 3 | 0.973 | 0.939 | 0.956 | 4914 |
| 4 | 0.874 | 0.908 | 0.891 | 878 |
| 5 | 0.854 | 0.945 | 0.897 | 629 |
| 6 | 0.828 | 0.896 | 0.861 | 366 |

| | | | | |
|---|---|---|---|---|
| 7 | 0.815 | 0.940 | 0.873 | 185 |

**Table S9A:** Performance of ESM-ECForest and comparison methods across the four levels of the EC hierarchy. Weighted F1, Micro F1, and Macro F1 scores are reported together with their 95% bootstrap confidence intervals (CI).

| Metric | EC level | Weighted F1 (95% CI) | Micro F1 (95% CI) | Macro F1 (95% CI) |
|---|---|---|---|---|
| ESM-ECForest | 1 | 0.944 [0.941, 0.947] | 0.943 [0.940, 0.946] | 0.911 [0.902, 0.919] |
| | 2 | 0.929 [0.925, 0.934] | 0.921 [0.917, 0.924] | 0.847 [0.837, 0.855] |
| | 3 | 0.920 [0.915, 0.924] | 0.912 [0.908, 0.916] | 0.826 [0.815, 0.835] |
| | 4 | 0.902 [0.896, 0.906] | 0.888 [0.883, 0.892] | 0.790 [0.778, 0.799] |
| DeepECTransformer | 1 | 0.925 [0.920, 0.930] | 0.925 [0.920, 0.930] | 0.889 [0.878, 0.900] |
| | 2 | 0.907 [0.901, 0.913] | 0.906 [0.900, 0.912] | 0.854 [0.841, 0.867] |
| | 3 | 0.889 [0.883, 0.895] | 0.887 [0.881, 0.893] | 0.815 [0.800, 0.830] |
| | 4 | 0.857 [0.850, 0.864] | 0.854 [0.847, 0.861] | 0.758 [0.740, 0.776] |
| ProteInfer | 1 | 0.918 [0.913, 0.923] | 0.919 [0.914, 0.923] | 0.863 [0.851, 0.875] |
| | 2 | 0.890 [0.884, 0.896] | 0.890 [0.884, 0.896] | 0.821 [0.807, 0.835] |
| | 3 | 0.873 [0.867, 0.879] | 0.871 [0.865, 0.877] | 0.783 [0.767, 0.799] |
| | 4 | 0.821 [0.813, 0.829] | 0.817 [0.809, 0.825] | 0.699 [0.678, 0.720] |
| DeepEC | 1 | 0.660 [0.652, 0.668] | 0.680 [0.672, 0.688] | 0.559 [0.541, 0.577] |
| | 2 | 0.631 [0.622, 0.640] | 0.644 [0.635, 0.653] | 0.511 [0.492, 0.530] |
| | 3 | 0.602 [0.592, 0.612] | 0.611 [0.601, 0.621] | 0.462 [0.442, 0.482] |
| | 4 | 0.548 [0.537, 0.559] | 0.553 [0.542, 0.564] | 0.389 [0.367, 0.411] |

**Table S9B:** Class-specific EC prediction performance of DeepECTransformer at EC Level 1.

| EC class | Precision | Recall | F1 score | Support |
|---|---|---|---|---|
| 1 | 0.948 | 0.936 | 0.942 | 4663 |
| 2 | 0.922 | 0.908 | 0.915 | 3212 |
| 3 | 0.941 | 0.929 | 0.935 | 4914 |
| 4 | 0.918 | 0.862 | 0.889 | 878 |
| 5 | 0.958 | 0.881 | 0.918 | 629 |
| 6 | 0.864 | 0.827 | 0.845 | 366 |
| 7 | 0.799 | 0.754 | 0.776 | 185 |

**Table S9C:** Class-specific EC prediction performance of ProteInfer at EC Level 1.

| EC class | Precision | Recall | F1 score | Support |
|---|---|---|---|---|
| 1 | 0.942 | 0.935 | 0.938 | 4663 |
| 2 | 0.915 | 0.894 | 0.904 | 3212 |
| 3 | 0.935 | 0.943 | 0.939 | 4914 |
| 4 | 0.890 | 0.841 | 0.865 | 878 |
| 5 | 0.904 | 0.849 | 0.876 | 629 |
| 6 | 0.829 | 0.799 | 0.814 | 366 |
| 7 | 0.733 | 0.673 | 0.702 | 185 |

**Table S9D:** Class-specific EC prediction performance of DeepEC at EC Level 1.

| EC class | Precision | Recall | F1 score | Support |
|---|---|---|---|---|
| 1 | 0.818 | 0.783 | 0.8 | 4663 |
| 2 | 0.762 | 0.729 | 0.745 | 3212 |
| 3 | 0.832 | 0.364 | 0.506 | 4914 |
| 4 | 0.683 | 0.618 | 0.649 | 878 |
| 5 | 0.718 | 0.632 | 0.672 | 629 |
| 6 | 0.586 | 0.502 | 0.541 | 366 |
| 7 | - | - | - | 185* |

*DeepEC does not provide EC 7 predictions. EC 7 was retained in the overall hierarchical evaluation as an unsupported class but is shown as N/A in class-specific evaluation.
